# Optimized iMAP CRISPR Perturbation Illuminates the Context-Specific Functions of RNA Modification Factors

**DOI:** 10.1101/2025.07.24.666689

**Authors:** Ge Wei, Zhengyu Jing, Weikai Wang, Bo Liu, Shaoshuai Mao, Junjun Lv, Xiaoming Zhang, Dingyu Cao, Ruilin Sun, Aibing Liang, Tian Chi

## Abstract

iMAP is a large-scale CRISPR perturbation platform based on a germline-transmitted transgene comprising individually floxed, tandemly linked guides that can be inducibly expressed via Cre-mediated recombination. A key limitation was strong recombination bias favoring the proximal guides. Here, we effectively mitigated this bias by inactivating a recombination hotspot, yielding guide representation nearly as uniform as traditional lentiviral libraries. Applying this optimized platform, we systematically profiled 70 (52%) RNA modification factors across 46 mouse tissues, uncovering pervasive tissue-specific essentiality. Notably, iMAP displayed much higher essential gene recovery than prior lentiviral CRISPR screens, attributable to its stronger guide depletion and lower technical noise. Finally, in a tumor model, iMAP identified *Thg1l* as a repressor of NK cell TNFα production; its ortholog knockout enhanced human NK cell-mediated tumor killing, underscoring therapeutic potential. All data are available in an interactive iMAP database. This work advances the Perturbation Atlas, a foundational resource for functional genomics.

**Teaser:** A unique CRISPR perturbation tool has been upgraded and deployed to reveal functions of epigenetic regulators in mice.

## Introduction

Decades after sequencing of the human genome, the context-specific functions of ∼20000 protein-coding genes across hundreds of cell types in the human body in health and disease remain poorly defined. A pressing task in functional genomics research is to construct “Cell and Tissue Perturbation Atlas”, which involves systematic disruption of mammalian genome and deciphering of the resulting phenotypes at the cellular and tissue levels in a high-throughput manner in both human and animal models (*1*). Such an atlas can reveal causal relationship between genetic circuits and cellular phenotypes, thus complementing various observational atlases and omics studies, such as the Human Cell Atlas aimed at profiling gene expression patterns of all human cell types (*1–3*). Importantly, with increasing amounts of perturbation data, artificial intelligence and machine learning can be deployed to predict gene function, thus enabling a Perturbation Atlas to serve as a generative causal foundation model, which will greatly empower our understanding of human biology in health and disease (*1*).

Pooled CRISPR screening is a standard and powerful high-throughput method for exploring gene functions in cell culture (*4*) and in transplanted immune cells (*5*), but its direct application to the cells *in situ* in mice remains challenging due to inefficient/uneven delivery of viral sgRNA libraries to target cells in their complex native environment(*6–9*). As a result, *in situ* screening has only been reported sporadically for a few solid organs (*10–17*). To eradicate the delivery problem inherent in viral libraries, thus achieving systematic *in situ* perturbation of diverse tissues throughout the whole mice, we have combined CRISPR-Cas9 and Cre-LoxP technologies into a virus-free platform named inducible mosaic animal for perturbation (iMAP) (*18–20*). iMAP uses a novel germline-transmitted transgene comprising 100 individually floxed gRNA-coding units (“guides”) linked together in tandem (Figure 1A). The array of guides is placed under the control of a modified U6 promoter with a LoxP variant (Lox71) embedded. In mice carrying the transgene and also ubiquitously expressing CreER and Cas9, treatment of mice with Tamoxifen (TAM), which activates Cre, would trigger recombination between Lox71 and one of the downstream LoxP variant sites preceding each guide (LoxTC9), leading to the expression of all the guides in the mice, but only one of them per cell, thus converting the mice into mosaic organisms ready for phenotypic characterization. iMAP is advantageous over conventional pooled *in situ* CRISPR screens, in that CRISPR perturbation can be readily induced via simple TAM administration, allowing parallel analyses of diverse cell types and tissues throughout the mice, with the mouse lines recurrently accessible as they can be maintained, propagated and shared just as conventional transgenic mice. Importantly, while pooled CRISPR screens typically use multiple gRNAs per target gene, the nonorthodox “one guide/target” design proves acceptable for iMAP (see Discussion) (*18*). With a single iMAP mouse line, and using gRNA representation as the readout, we have rapidly constructed a miniature version of the mammalian perturbation atlas cataloguing the effects of perturbations of 90 genes across 39 organs/tissues/cells (“tissues” hereafter), which is hardly achievable using other tools. However, a major weakness of iMAP is that the recombination between Lox71 and the series of downstream LoxTC9 sites is biased toward the proximal (mainly the first 11 or g1-11) sites, leading to the underrepresentation of the distal guides at P0 in the post-recombination library, which necessitates the use of more cells in order to cover the distal guides (Figure 1A) (*18*). The bias in recombination thus compromises assay sensitivity, hampering the profiling of rare cells in the body.

**Figure 1.**
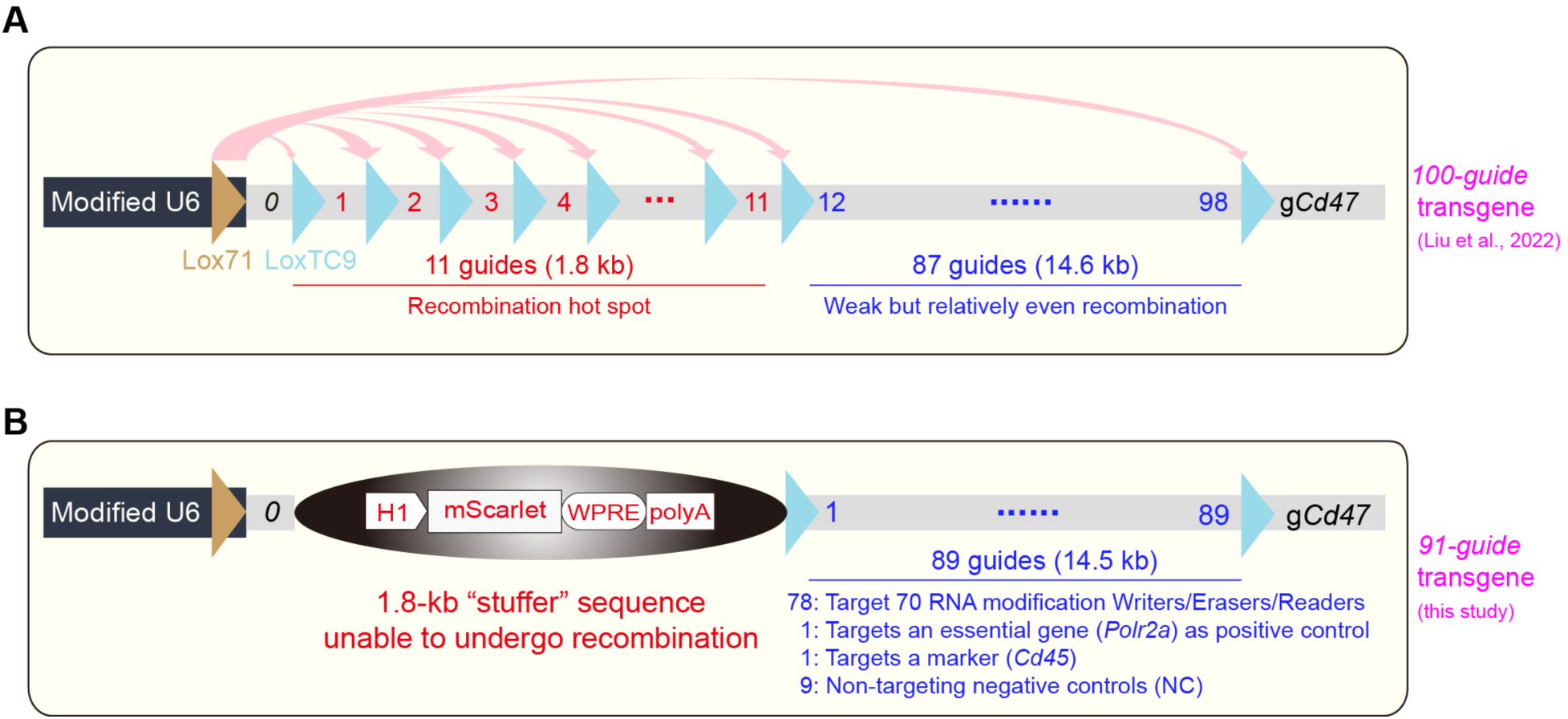
Strategy for minimizing recombination bias. (A) First-generation iMAP transgene. Lox71 and TC9 are LoxP variants embedded in the U6 promoter and precede each gRNA-coding gene (“guide”), respectively. The guide at Position 0 (P0) is g*Cd19* (not shown). The last guide (g*Cd47*) facilitates PCR-NGS quantification of recombined products. Following recombination between Lox71 and a LoxTC9 preceding a guide, the intervening sequence is deleted, re-locating the guide to P0 for transcription. The recombination between Lox71 and the series of LoxTC9 sites is biased toward the proximal LoxTC9 sites except the first site, perhaps due to steric hindrance created by the transcription machinery at U6 promoter. (B) Optimized configuration. The “stuffer” harbors an expression cassette for monitoring U6 promoter activity, which proved unsuccessful (Figure S1D). The guide at Position 0 (P0) is g*Cd19* (not shown). The target genes and gRNAs are listed in Table S1.

All known types of RNA, including mRNA, tRNA and rRNA, are subject to dynamic chemical modifications, which impacts RNA stability, subcellular localization and/or interactions with other biomolecules (*21–23*). The modifications are deposited and removed by the “writers” and “erasers”, respectively, whereas the “readers” bind the modification marks and recruit effector machinery to alter cell functions, thus translating the modification signals into functional outputs. RNA modification has emerged as a crucial regulator of gene expression in health and disease, and is attracting increasing attention. The latest version of the Modomics Database documents 335 RNA modifications in nature and 137 modification factors (writers, erasers and readers) in humans (https://genesilico.pl/modomics/) (*24*). Traditionally, RNA modification is investigated mostly in cell culture. KO mice have also been described for some of the modification factors (see further), but the KO often proves lethal, precluding in-depth characterization. Thus, the roles of the modification factors in mice, especially in various cell types in adults, remain largely unexplored.

In this study, we substantially improved the uniformity of guide representation in iMAP, which is now close to that in conventional lentiviral sgRNA libraries used in pooled CRISPR screens. We then interrogated the functions of 70 RNA modification factors across 46 organs/tissues/cell types (“tissues” hereafter) in adult mice. Furthermore, we combined iMAP and a tumor model to identify a candidate gene for potentiating tumor-killing function of NK cells, which was subsequently validated in human NK cells. Finally, we describe a public iMAP database.

## Results

### iMAP optimization via elimination of recombination hotspot

The Lox71-TC9 recombination in the first-generation iMAP transgene is biased toward roughly the first 11 guides (g1-11), indicating that the region encompassing ∼g1-11, about 1.8 kb in length, is a recombination “hot spot” (Figure 1A). We reasoned that its replacement with a 1.8-kb inert “stuffer” sequence lacking LoxP sites might force Lox71 to skip the “hotspot” to reach out to and rearrange with the downstream TC9 sites, thus mitigating the bias in recombination. To test this, we constructed a *91-guide* transgene with a 1.8-kb stuffer inserted between g0 and g1 (Figure 1B; see further for guide composition). In the ensuing transgenic mice, the U6 promoter was active, unmethylated and the *91-guide* array intact (Figure S1A-C). We profiled P0 guides in adult *91-guide; Cre* mice at various levels of TAM-induced recombination, and compared the profiles with that in *100-guide; Cre* mice at similar levels of recombination. There is indeed substantial improvement in recombination uniformity in the *91-guide* array at all levels of recombination (Figure 2A), which is more obvious if the guides are analyzed as groups (Figure 2B-C). Specifically, for the *100-guide* array, the group comprising g1-10 was predominant, constituting up to 49-72% of the total recombined P0 guides (yellow bar, Figure 2C top), contrary to the distal groups especially g41-99 (1-5%; last six bars, Figure 2C top). In contrast, in *91-guide; Cre* mice, g1-10 dropped to 26-34% while g41-90 rose to 6-15% (yellow vs last five bars, Figure 2C bottom), indicating marked reduction in recombination bias.

**Figure 2.**
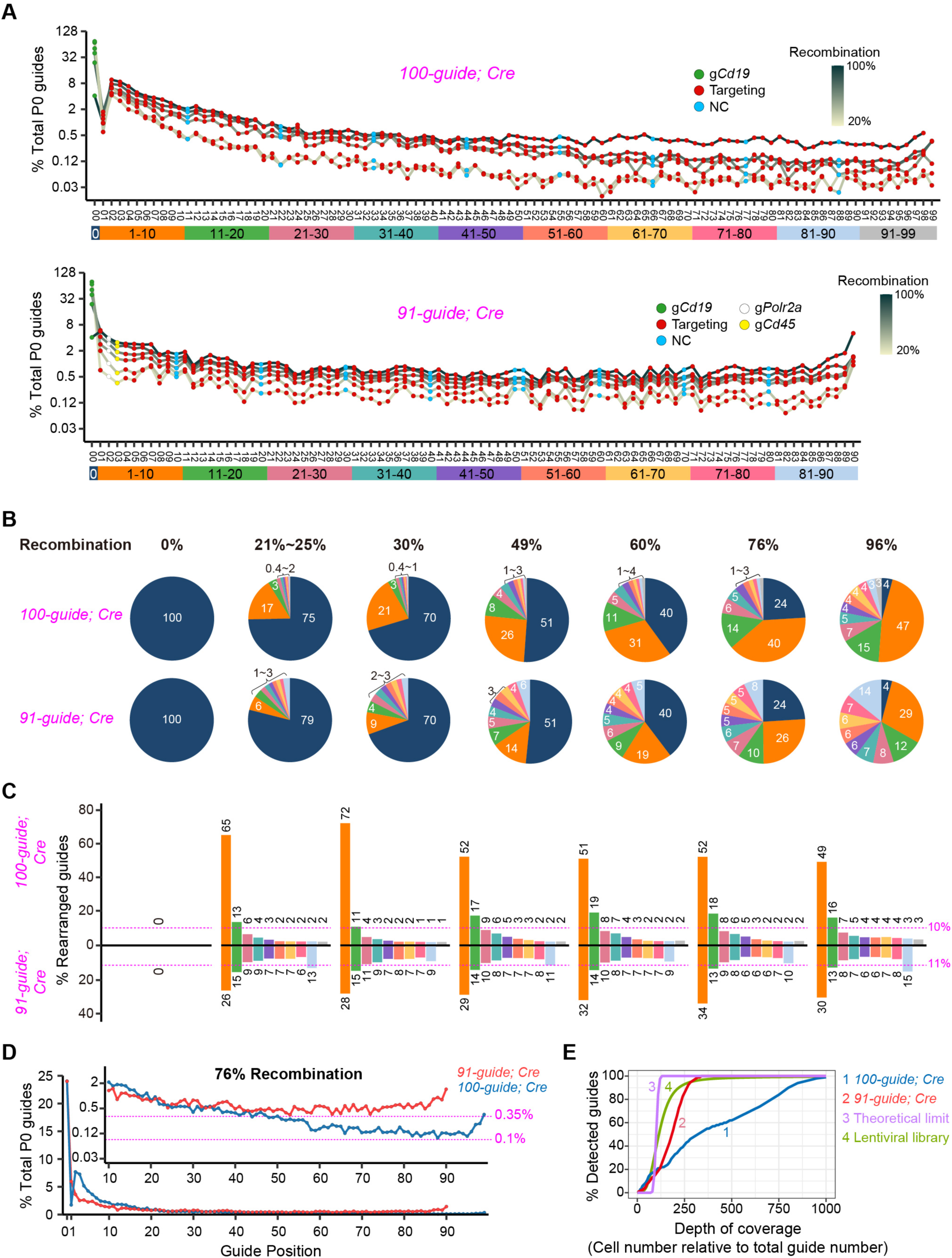
Improved guide representation in the optimized iMAP transgene. (A) P0 guide profiles as a function of recombination extents. *100-guide; Cre* (top) or *91-guide; Cre* (bottom) mice were treated with increasing doses of TAM before analysis of various organs (Figure S1E), and samples with progressive recombination, as measured by g*Cd19* deletion, are selected for display. Shown is the relative abundance of individual guide among all P0 guides, with each line representing a sample. The guides are arranged according to their positions on the array. An ID number is appended to each guide if several guides are designed for the same genes. Two identical non-targeting guides (g50, 51) were inadvertently built into the *91-guide* array. The colored bar at the bottom specifies the groups of guides whose abundance is analyzed in B. Source data in Table S2. (B) Same as A, but displays the guides as groups defined by the colored bar in A. The values above the pies indicate the extent of g*Cd19* deletion; e.g., the third pie shows 70% of g*Cd19* survived recombination, translating to 30% deletion. The upper and lower pie charts are paired. (C) Same as B but displays guide abundance relative to all the re-arranged P0 guides (hence with g*Cd19* excluded), which better exposes the bias in recombination. The dotted lines denote the theoretical guide abundance if the recombination is absolutely even. (D) Same as A, but compares a pair of samples with 76% recombination from the two mouse lines. The inset displays the distal and hence low-abundance guides. (E) Monte Carlo simulation of guide detectability as a function of number of the cells sampled. The simulation is based on actual guide representation after recombination (#1-2) or viral transduction (#4; data averaged from two lentiviral libraries reported in (*54*).)

iMAP sensitivity is limited by the least abundant P0 guide generated after recombination. We found that in representative samples from the two mouse lines, each with 76% recombination (a level typically achieved in our routine experiments), such guides constituted 0.1% and 0.35% of the total P0 guides for the *100-* and *91-guide* transgene, respectively (Figure 2D). Thus, the stuffer led to ∼3.5x increase in sensitivity.

For iMAP analysis of rare cells, it can be challenging to isolate enough cells to cover all the guides in the library, particularly the underrepresented guides. We thus used mathematical modeling to explore the relationship between the numbers of cells sampled and the % of the guides detected. We assume a detected guide is represented by at least 100 cells, a depth we previously found sufficient for iMAP analysis, and we express cell number in relative terms, as “fold over the total guide number in the array” (e.g., a value of 1 for the *100-guide* array means 100 cells sampled). For *100-guide*, the model predicts that the cell number must be at least 1000 times the guide number (namely 100,000 in absolute number) to detect ∼100% of the guides, and the value drops to 60% and 40% when the cell number drops 2x and 4x, respectively (Figure 2E, line 1). In contrast, for *91-guide*, cells only 300 times the guide number suffice to cover nearly all the guides, reinforcing the notion that the stuffer led to 3x increase in iMAP sensitivity (Figure 2E, line 2). This sensitivity is still 3x lower than the theoretical limit achieved if the recombination becomes absolutely even (Figure 2E, line 3), but already surpasses that of conventional lentiviral library, in that full coverage of the viral library requires cells 500 times the gRNA number, contrary to only 300 times for *91-guide* (Figure 2E, line 4).

However, for partial coverage, the viral library is more sensitive. For example, to cover 80% of the libraries, pooled CRISPR screen and iMAP would need 170x and 240x cells, respectively (Figure 2E, line 4 vs 2). Thus, on balance, the two platforms perform similarly. Of note, for pooled CRISPR screen, Unique Molecular Identifier (UMI) can be used to improve the screen sensitivity, but this strategy is not applicable to iMAP, which is a limitation of the latter platform.

We conclude that the stuffer led to 3.5x increase in iMAP sensitivity, making it comparable to that of the classic lentiviral sgRNA library.

### Overview of iMAP-91 phenotype

As depicted in Figure 1B, the *91-guide* transgene features 78 guides targeting 70 RNA modification genes (writers/erasers/readers), each gene targeted by one guide except that five genes (*Alkbh3, Fto, Mettl3, Ythdc1, Ythdf1*) are each covered by several guides (to verify the robustness of the “one-guide/target design”; see Discussion). The array also carries a positive control targeting *Polr2a*, an essential gene whose perturbation causes cell death, and nine non-targeting guides (negative control, NC) spiked into the array to control for bias in recombination. We bred *91-guide; Ubc-CreER; CAG-Cas9* mice (“iMAP-91”), treated them with TAM around D10 and analyzed guide representation in diverse tissues 3-4 months thereafter in three females and three males. The mice were grossly normal except for atrophic testes (Figure 3A). We profiled P0 guides in 46 tissues sampled from six mice, yielding 4,002 data points (87 important guides x46 tissues) each averaged from at least two (mostly 5-6) mice, the replicates being highly reproducible (Figure S2). Guide depletion events were predominant, constituting 49% of the 4,002 data points using fold ≥ 2 as the threshold (black lines) or 74% using gNC for comparison (green lines), contrary to only 0.1%–3% for the enrichment and 23%–50% for the unaltered (Figure 3B). Note that the gNCs formed a sharp peak, attesting to the low noise level of the system. We used simple bar graphs to display the enriched guides (Figure 3C), and a heatmap to display depletion where the enrichment is scored as “unaltered” to better visualize the depletion (Figure 3D). Guide enrichment was marginal (mostly 1.5x) and highly restricted, detected for only four guides (g63-*Rpusd4,* 15*-Trdmt1,* 48*-Snu13,* 62*-Trmu*) and in only two organs (eyeball and cerebellum; Figure 3C); KO data for these genes are scarce but not in conflict with the iMAP data (see Discussion). In contrast, guide depletion was pervasive, encompassing many genes with established functions, thereby enabling rigorous assessment of the depletion’s biological validity, as detailed below.

**Figure 3.**
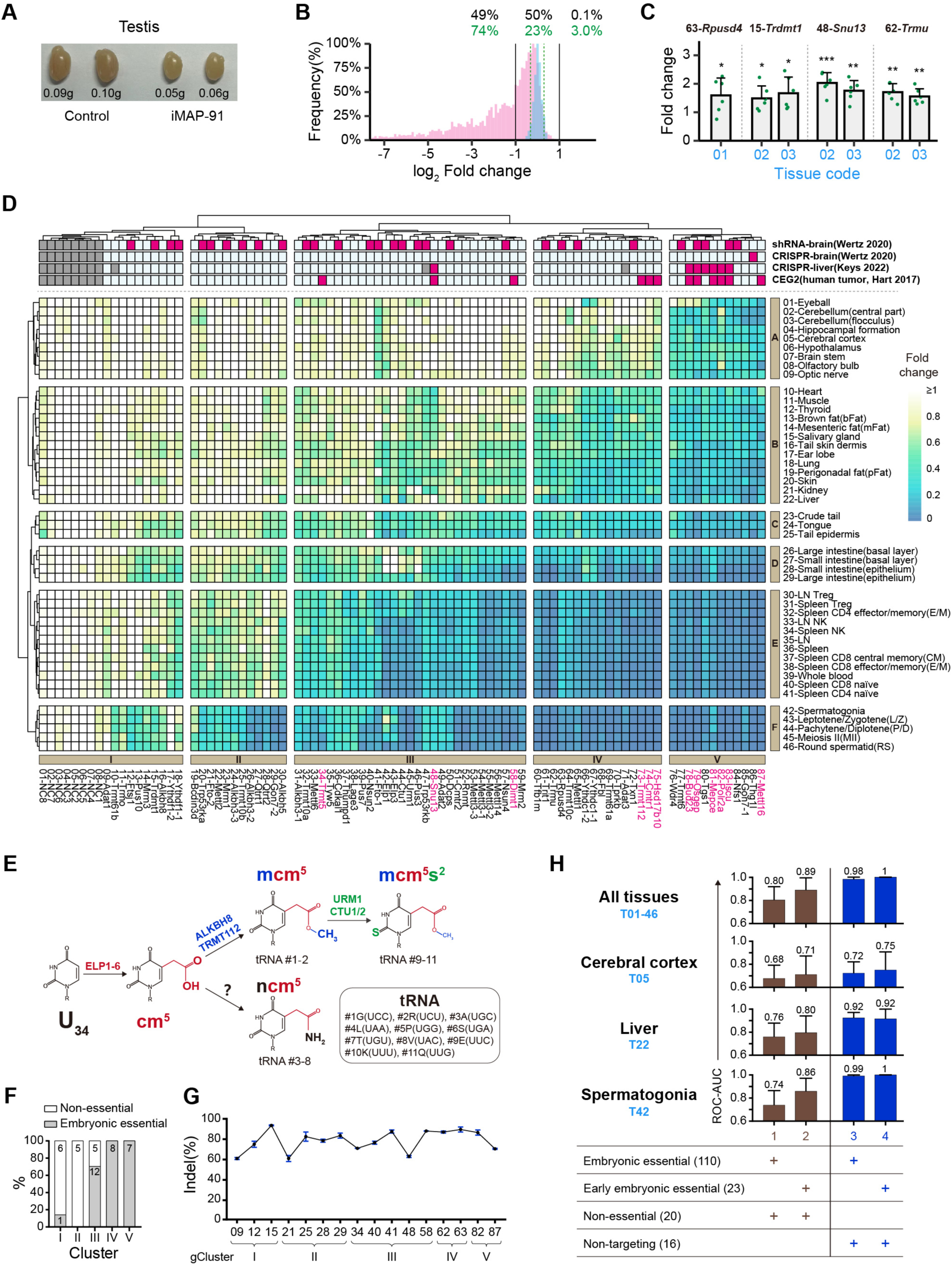
iMAP-91 characterization. (A) Atrophic testes. (B) Frequency distribution of the fold-changes for the 79 targeting (pink) and eight NC (cyan) guides across 46 tissues. The 4002 (87×46) datapoints were binned according to the fold-change magnitudes, and the frequencies of each bin displayed for each guide, with the highest frequency in each histogram set as 100%. Black lines mark 2-fold changes, whereas the green lines indicate the 5th and 95th percentiles of gNC values. (C) Guide enrichment. Tissue code 01, 02, 03 refers to eyeball, cerebellar central part and cerebellar flocculus, respectively, as specified in (D). Values are mean ± SD (n=6). *p < 0.05, **p < 0.01, ***p< 0.001, Student’s t-test. (D) Cluster analysis of guide depletion. Displayed are 87 guides comprising 78 guides targeting RNA modification factors, one targeting *Polr2a* (g82), and eight gNC (g1-8). Gene essentiality per literature is indicated at the top of the heatmap, where shRNA-brain and CRISPR-brain refer to, respectively, the shRNA- and sgRNA-based genomewide *in situ* screens for neuronal essential genes in the mouse striatum (*16*), CRISPR-liver is a sub-genomewide *in situ* screen for liver essential genes (*11*), and CEG2 (pink) the core essential gene set identified in human tumor cell lines(*31*); three genes targeted by iMAP-91 are not covered in the CRISPR-liver screen (gray box). Each data point (guide-tissue combination) is averaged from 2 to 6 (mostly 5-6) biological replicates. Source data in Table S3. (E) Cascade reaction leading to tRNA-U34 modifications (*46*). (F) Guide clustering coincides with the phenotypes of global single-gene KO mice. Analyzed are the genes whose KO phenotypes are available from MGI. The number of genes in each category is indicated. *Alkbh3* fits both Cluster II and III because it has three guides, two (g24, 26) residing in Cluster II and the slightly distant third (g31) at the very beginning of Cluster III. Source data in Table S3. (G) Editing rates of representative gRNAs. gRNAs and Cas9 were co-expressed in N2a cells in duplicates before indel quantification. (H) ROC-AUC analysis. The data are pooled from iMAP-91 and iMAP-100. The numbers of targeted genes and non-targeting guides are bracketed. Error bars represent 95% confidence intervals (CI, Delong’s method). The analysis was performed using pROC in R. The KO phenotypes are compiled from MGI (https://www.informatics.jax.org/). Source data in Table S3.

### Guide depletion pattern is biologically relevant

In the guide-depletion heatmap, genes clustered into five groups (I–V) and tissues into six (A–F; Figure 3D). We previously identified four salient features in the iMAP-100 heatmap that strongly support the biological relevance of this clustering (*18*). These features are all recapitulated in iMAP-91, as detailed below.

First, related guides tended to co-segregate. Thus, the eight gNC tightly co-segregated (gCluster I), as were guides targeting different essential subunits of the same protein complexes, including the following complexes: *Mettl3-Mettl14* (g53-56), *Iscu-Nfs1* (g83-84), *Qtrt1-Qtr2* (g27-28) and to some extents *Trmt6-Trmt61a* (g77,69) and *Mettl1-Wdr4* (g65-76) (Figure 3D)(*25–30*). Besides, the guides targeting four genes required for tRNA-U34 modifications (*Elp1*, *Elp3*, *Ctu1*, *Urm1*) tightly co-segregated (g42-45), although the guides targeting two other genes in the pathway (g16-*Alkbh8* and g73-*Trmt112*) did not (Figure 3E; see Discussion). Finally, different guides designed to inactivate the same genes, namely *Mettl3* (g53-55), *Ythdc1* (g66-67), *Ythdf1*(g17-18) and *Alkbh3* (g24,26,31), all (tightly) clustered (Figure S3A). On the other hand, guides predicted to impact the same genes but to different extents did not co-segregate, as in the case of *Fto* (g21, 41) and *Gon7* (g29, 85), albeit with the caveat that g85-*Gon7* additionally targets two unannotated genes related to *Gon7* (Figure S3B-C).

Second, relevant tissues tended to co-segregate just as relevant guides, including neural tissues (T2–9), skin-related tissues (T16-17, 20, 23, 25), intestine-related tissues (T26–29), lymphocyte subsets (T30–41), and male germ cell subsets (T42–46). Of note, different tissues apparently have different sensitivities to Cas9, with post-mitotic tissues relatively resistant, which poses a challenge for Perturb Atlas mapping (see Discussion).

Third, the genes generally required for cell survival in iMAP-91 were enriched with the ‘‘core essential genes’’ (CEGs) generally required for human tumor cell survival/proliferation (*31*). Specifically, of the 71 mouse genes targeted in iMAP-91, 12 were orthologous to human CEGs, all distributed among the last 15 mouse genes (73–87) in the heatmap except three (34-*Trmt5*, 48-*Snu13*, 58-*Dimt1* (Figure 3D, pink text); the exceptions have medical implications (see Discussion).

Fourth, guide depletion patterns in iMAP mice mirrored the phenotypes of conventional global single-gene KO mice. Thus, ablation of gCluster IV-V genes is all embryonic lethal (15/15), gClustre III mostly so (12/17) whereas gCluster I-II all viable except for one gene (1/12; Figure 3 F; Table S3).

The following analyses further affirm the heatmap’s biological relevance while highlighting iMAP’s superior performance relative to conventional CRISPR screens.

First, we measured editing efficiencies for 16 guides—sampled to represent all clusters in the heatmap—in a mouse tumor cell line. All were active (60–90% indel rates), with no correlation between indel efficiency and cluster assignment, confirming that clustering reflects gene functions rather than guide potency (Figure 3G).

Second, we computed the receiver operating characteristic area under the curve (ROC-AUC), the standard metric for negative-selection CRISPR screens that measures how well validated essential genes rank as more depleted than non-essential genes (*32*, *33*). Lacking gold-standard lists for primary mouse cells, we used organismal viability as a proxy. To enhance statistical power, we pooled data from iMAP-91 (71 genes; Figure 3D heatmap) and iMAP-100 (90 genes; Figure 4D heatmap) (*18*), yielding 161 genes total. MGI single-gene knockout data classified 110 as embryonic essential (EE; lethal before E21), including 23 early embryonic essential (EEE; lethal between E4.5–E8, more likely cell-essential), and 20 as non-essential (no lifespan shortening; Table S3). Using non-essential genes as negatives and averaging log₂ fold changes across all surveyed tissues, we obtained ROC-AUC of 0.80 (95% CI: 0.69–0.92) or 0.89 (95% CI: 0.78–0.99) with EE or EEE as positives, respectively (Figure 3H, top plot, columns 1–2). Strikingly, ROC-AUC rose to 0.98–1.00 when non-targeting guides defined the negative class, indicating minimal technical noise in iMAP (Figure 3H, top plot, columns 3–4). Tissue-specific analyses (cerebral cortex, liver, and spermatogonia, representing low, intermediate, and high Cas9 sensitivity, respectively) confirmed that ROC-AUC increased with sensitivity (e.g., 0.71, 0.80, 0.86 using EEE positives and non-essential negatives; Figure 3H, column 2). Notably, these values surpass those from a lentiviral screen (see Discussion).

**Figure 4.**
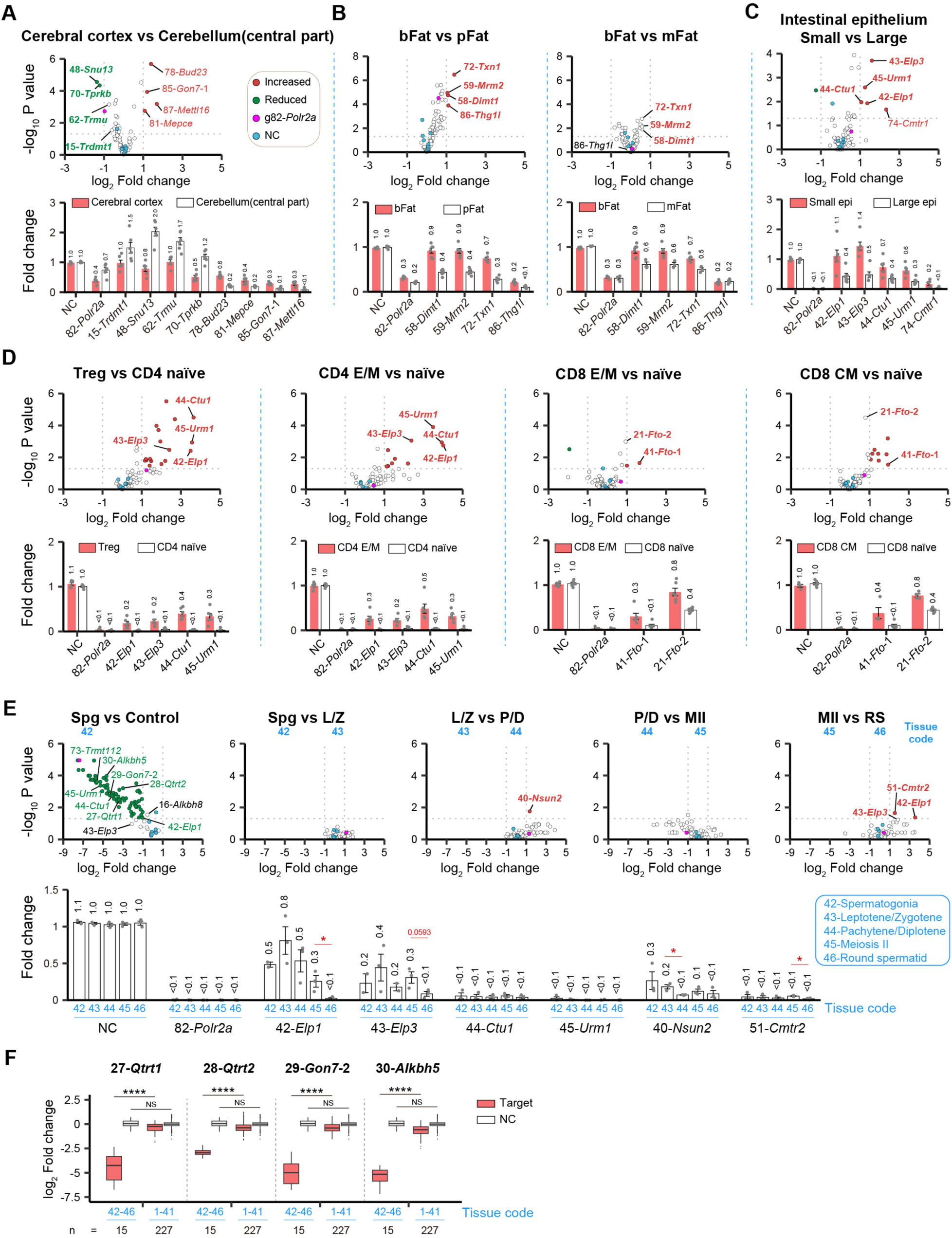
Mining the heatmap data. (A-D) Comparing tissue pairs. Shown are differentially represented guides (top) and their absolute abundance normalized to *91-guide; Cre* control mice (bottom). Red and green dots within the top plots indicate significant enrichment and depletion, respectively (fold-change> 2, p<0.05). (E) Guide depletion during spermatogenesis. Same as A-D, but focusing on the male germline. (F) Germline-specific guide depletion. Values are mean ± SD, n = 15–227, each data point being a guide-tissue-replicate combination. ****p < 0.0001.

Finally, comparisons with genome-wide lentiviral screens in brain(*16*) and liver (*11*) revealed that iMAP-91 uncovered far more essential genes: 12 core essentials (gCluster V) and 16 liver essentials (gCluster IV), versus only one of these in brain and six in liver from the viral screens (marked atop the heatmap; Figure 3D).

Collectively, these data underscore iMAP’s outstanding performance. We next mined the heatmap as described below.

### Tissue Cluster A: differential effects of CEGs and uniqueness of cerebellum

In this group of tissues (brain, heart, and eyeball), guide depletion was undetectable/weak except for gCluster V (g76-87), which was clearly depleted in all tissues, suggesting that g76-87 had targeted CEGs. Guide alteration patterns were similar among various brain regions, but comparison of cerebral cortex vs cerebellar central part (Tissue 05 vs 02) revealed multiple guides with differential representations, which occurred due to either quantitative (g78, 81, 82, 85, 87) or qualitative (g15, 48, 62, 70) differences in guide abundance between the two tissues (Figure 4A, top). Specifically, in the “quantitative” group, the guides were depleted in both tissues but to different extents, with g82 preferentially depleted in the cerebrum while the remaining (g78, 81, 85, 87) just the opposite. Of note, these five guides (g78, 81, 82, 85, 87) all target CEGs, suggesting that although CEGs are essential for cell survival, the kinetics of cell death following CEG KO could be cell-type dependent. This said, for the most part, gCEGs are depleted to similar extents in closely related tissue pairs and thus are useful controls for Cas9 cutting efficiencies (see, e.g, g82-*Polr2a* in Figure 4B-E). Regarding the “qualitative” group, three guides (g48, 15 and 62) were selectively enriched in the cerebellum but no other tissue, whereas g70 generally depleted in various tissues except cerebellum, thus exposing a unique aspect of the cerebellum (Figure 4A and 3D).

### Tissue Cluster B: brown vs white fat

Brown fat (bFAT) burns calories and is located primarily in the interscapular region, whereas white fat stores excess energy and is distributed widely, with perigonadal fat (pFAT) and mesenteric fat (mFAT) among the major white adipose tissues. Three guides (g58, 59, 72) were selectively depleted in pFAT and mFAT but not bFAT, suggesting that their target genes (*Dimt1*, *Mrm2* and *Txn1*) selectively promote white adipocyte survival/proliferation/development without impacting bFAT (Figure 4B).

### Tissue Cluster D: small vs large intestine

Five guides were differentially represented in the epithelium of small vs large intestine, which remarkably include the “tetra-guides” targeting tRNA-U34 modification (g42-*Elp1*, 43-*Elp3*, 44-*Ctu1*, 45-*Urm1*; Figure 4C). Specifically, all the four guides were depleted clearly (2-4x) in the epithelium of large intestine, whereas in the small intestine epithelium, the depletion was marginal (∼1.5x) and restricted to g*Ctu1/Urm1*, suggesting preferential requirement of tRNA-U34 modification for the large intestine.

### Tissue Cluster E: Regulators of T cell development and activation

gCluster III–V guides were generally depleted across various immune cells but occasionally to different extents in pairs of closely related T cell subsets: First, over a dozen guides were preferentially depleted in naïve CD4 T cells relative to Tregs, with the tetra-guides that target tRNA-U34 modification among the top candidates (Figure 4D, panel 1). The two T cell subsets develop from the same precursor (DP thymocytes), suggesting that this lineage bifurcation is impacted by tRNA-U34 modification.

Second, the tetra-guides targeting tRNA-U34 modification were also the top preferentially depleted candidates in naïve relative to effector/memory(E/M) CD4 T cells (Figure 4D, panel 2). Thus, tRNA-U34 modification seems a prominent repressor of both Treg development and CD4 T cell activation/differentiation. This function is specific, being absent from CD8 T cells; instead, FTO, which erases N^6^-methyladenosine (m^6^A), emerged as a prominent repressor of CD8 differentiation into both E/M and central memory (CM) CD8 T cells (Figure 4D, panel 3-4). Of note, multiple other guides were also enriched in CM relative to naïve CD8 cells (Figure 4D, panel 4, red dots in the top plot), but these guides were severely depleted even in CM CD8 cells, making their relative enrichment in CM CD8 cells less interesting (not shown).

### Tissue cluster F: Spermatogonia vulnerability

Our previous analysis of iMAP-100 mice reveals striking vulnerability of spermatogonia, which is recapitulated in iMAP-91: First, spermatogonia were more vulnerable than their progeny (Figure 4E): out of the 79 targeting guides, only seven failed to impact spermatogenesis; the remaining 72 guides were depleted at least 2x in spermatogonia, and no further change was seen in the progeny except for g*Nsun2, Cmtr2* and *Elp1* (and to some extent g*Elp3*) whose depletion emerged in spermatogonia and exacerbated in P/D (for g*Nsun2*) or RS (for g*Elp1/Elp3/Cmtr2*; Figure 4E). In other words, 72 guides blocked spermatogonia survival/proliferation/renewal, but only three of them additionally impacted its progeny. Of note, for the four guides targeting tRNA-U34 modification, depletion in spermatogonia was moderate (2-4x) for g*Elp1/3* but dramatic (17-38x) for g*Urm1/Ctu1*, highlighting their functional dichotomy (see Discussion).

Second, spermatogonia were more vulnerable than somatic cells, in that gCluster II was preferentially depleted in spermatogonia relative to somatic cells (Figure 3D). In particular, four guides (27-*Qtrt1*, 28-*Qtrt2*, 29-*Gon7-2* and 30-*Alkbh5*) were depleted marginally in somatic cells but severely (6-27x) in the germline (Figure 4F). QTRT1 and QTRT2 form the heterodimeric tRNA guanine transglycosylase (TGT) that catalyzes the base-exchange of a wobble guanine (G34) with queuine (Q) in some tRNAs, with QTRT1 being the catalytic subunit(*34*). The co-depletion of both *Qtrt* guides strongly argues for the essential and selective role of TGT in spermatogenesis. Regarding *Gon7*, it encodes a subunit of the KEOPS complex that introduces tRNA N6-threonylcarbamoyladenosine (t6A) modification(*35*). g29-*Gon7-2* is predicted to eliminate the C-terminal 13 residues of GON7, thus revealing the selective and essential role of this tail in spermatogonia (Figure S3C). Finally, *Alkbh5* encodes an m6A eraser, and its global KO induces germ cell apoptosis and male sterility, but the mice seem otherwise normal, in general agreement with the iMAP data (*36*).

In summary, these data highlight the vulnerability of spermatogonia, which has medical implications (see Discussion).

### Uncovering a potential target for boosting human NK cancer therapy

Finally, we sought to enhance human cancer immune therapy, focusing on natural killer (NK) cells, a pivotal first-line tumor killer (*37*, *38*). Release of TNFα is a key mechanism of NK-mediated killing (*39*). Our strategy was to first use iMAP to identify potential candidate genes whose KO could elevate TNFα expression in mouse NK cells, and then use the information to engineer human NK cells, which simultaneously serves to validate the candidates; a similar strategy has been taken to potentiate human CAR T cells in our previous study (*18*).

Specifically, we inoculated iMAP-91 mice with mouse colon cancer MC38 cells, administered TAM to induce recombination, isolated and compared TNFα^hi^ vs TNFα^low/-^ (Figure 5A). A single guide (g86-*Thg1l*) was markedly enriched in the TNFα^hi^ relative to the TNFα^low/-^ subset (Figure 5A), implicating *Thg1l*, a probable tRNA^His^ guanylyltransferase, as a repressor of TNFα expression in mouse NK cells. To validate and translate this finding, we generated *THG1L* knockout human NK cells. Upon co-culture with K562 leukemia targets, *THG1L* KO NK cells secreted more TNFα (481 vs. 337 pg/mL) and exhibited enhanced target-cell lysis (65% vs. 39%) (Figure 5 B-C). Notably, expression of perforin, granzyme B, and the degranulation marker CD107a remained unchanged (Figure 5C), indicating that increased TNFα drives the improved killing. The phenotype was reproducible across biological replicates and with independent guide RNAs (Figure S4). This proof-of-concept mirrors our prior success in engineering CAR T cells (Liu et al., 2022) and establishes iMAP as a rapid platform for discovering immunomodulatory targets translatable to human cell therapy.

**Figure 5.**
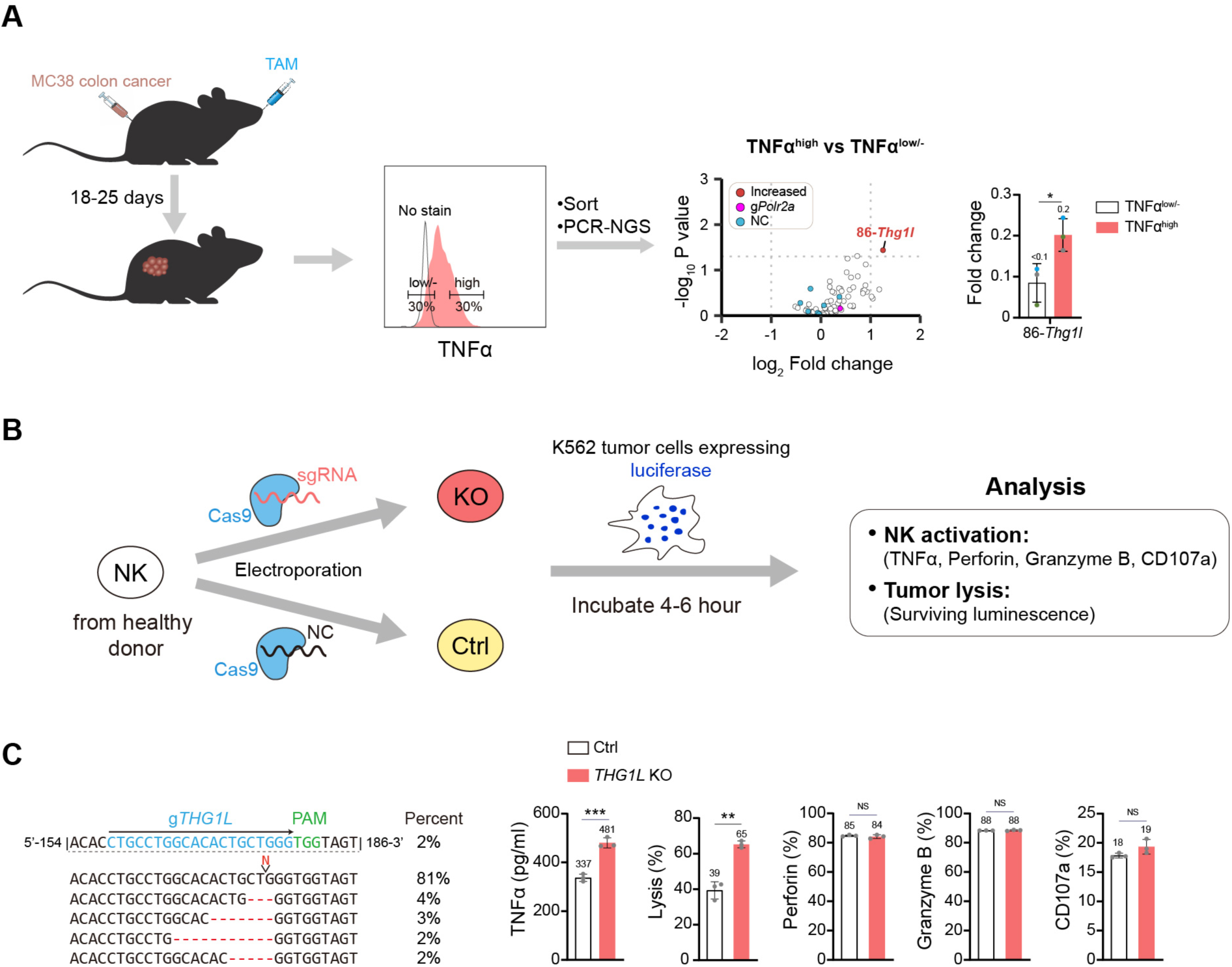
Screening and validation of NK regulators. (A) Genetic screens for NK regulators in tumor-bearing iMAP-91 mice. Tumor-infiltrating cells were stained for CD45.2, CD3 and NK1.1 together with intracellular TNFα before indicated NK subsets were isolated by flow cytometry and analyzed by NGS. Each data point in the volcano plot is averaged from 3 biological replicates. The bar graph displays the raw data (fold-changes) for g*Thg1l*, where the three biological replicates are indicated by dots with different colors. *p < 0.05, Student’s t-test. Source data in Table S3. (B) Workflow for target validation in human NK cells *in vitro*. (C) Successful editing and its effects on NK function for *THG1L*. Data are from one representative experiment, with a biological replicate shown in Figure S4A. *THG1L* KO human NK cells were incubated with K562 tumor cells for 4-6 hr before FACS analysis. The data show that *THG1L* KO increased TNFα secretion in NK cells and enhanced NK cells mediated lysis, but did not affect human NK degranulation, perforin or granzyme B. Dots represent technical repeats (n=3). *p < 0.05, **p < 0.01, ***p< 0.001, ****p < 0.0001, Student’s t-test.

### iMAP website

We have constructed a public database for browsing, searching and analyzing iMAP data (http://49.235.152.12/; Figure S5A). Four key modules are featured: “Heatmap”, which displays interactive *Guide x Tissue* atlases (Figure S5B); “Gene Search”, which enables query for the abundance of guides of interest across tissues (Figure S5C); “Network”, which presents an interactive network of the guides surrounding a guide of interest, with relevant information indicated, such as fold-changes of the guides, the number of tissues impacted by a guide, the distances among the guides and the total number of guides connected with a particular guide (Figure S5D); and “Tissue comparison”, which identifies the guides differentially represented between a pair of selected tissues (Figure S5E). Future efforts should culminate in the creation of the panoramic Perturbation Atlas covering all the genes across all the cell types at tissue and single-cell levels.

## Discussion

### Perturbation Atlas: challenges and countermeasures

A central goal in functional genomics is to build a comprehensive Perturbation Atlas that maps the *in vivo* phenotypic consequences of gene knockout across diverse tissues. This tissue diversity poses a challenge to iMAP (and classic CRISPR screens), because Cas9 cleavage efficiency varies markedly between tissues. Specifically, post-mitotic tissues (especially the brain) are relatively refractory to Cas9 probably due to limited chromatin accessibility; a potential solution here is to use Cas13 (which acts on mRNA) for perturbation. On the other hand, the male germline was highly sensitive to Cas9, with wide-spread guide depletion not only in embryonic essential genes but also non-essential genes. This extraordinary depletion pattern unlikely reflects cell death resulting from Cas9 induced DNA breaks. First, non-essential gene depletion was much weaker, enabling their clear separation from early essential genes (ROC-AUC = 0.86). More importantly, in an iMAP line targeting 6 marker genes each with 10 guides (iMAP-61), testicular guide abundance perfectly reflects known biology: depletion of cell-essential controls (*Polr2a, Kpnb1*), enrichment of tumor suppressors (*Pten, Trp53*), and no change in lineage markers not expressed in testis (*Cd19, Cd45*) (*18*). This, the widespread depletion in the male germline reflects genuine biological requirements. Anyway, given the variability in Cas9 cleavage, direct comparisons of guide abundance across distantly related tissues (e.g., brain vs. testis) are not meaningful. However, comparisons within closely matched tissue pairs of similar cellular state and Cas9 sensitivity—such as cerebellum vs. cerebrum or male germ cells at different developmental stages —remain valid and informative (Figure 4).

### iMAP robustness despite one guide per target

Although conventional CRISPR libraries employ multiple guides per gene to reduce false hits, iMAP deliberately uses only one guide to maximize library capacity, which fortunately does not markedly sacrifice robustness thanks to several design features. First, iMAP guides were selected using the state-of-the-art Rule Set 2 algorithm—the same algorithm that powers the Brunello library. In tumor cell viability screens, a single Brunello guide per gene already achieves excellent performance (ROC-AUC = 0.93), with additional guides providing only minor improvement (up to 0.98 with 4 guides/target)—in sharp contrast to the earlier GeCKOv2 library, which requires 6 guides to increase ROC-AUC from 0.73 to 0.91 (*32*). The iMAP libraries were designed according to the same principles as Brunello and therefore deliver comparable reliability with just 1 guide/target. Second, iMAP founder lines were pre-screened by RT-qPCR for strong gRNA expression. Third, all guides are integrated at a single genomic locus in every cell, minimizing cell-to-cell variability. Finally, iMAP exhibits exceptionally low technical noise (see further). These features collectively deliver impressive performance across a series of benchmarks: false hits are rare (1/10 false negatives and 1/590 false positives in the iMAP-61 control line targeting marker genes) (*18*); iMAP phenotypes consistently recapitulate known biology; all 16 guides randomly selected from iMAP-91 proved active (Fig. 3G); and finally, as elaborated in the next three sections, distinct guides targeting the same genes elicit similar phenotypes, seeming inconsistencies with the literature are reconcilable, and iMAP performance appears to rival that of classic CRISPR screens.

### Consistencies among orthogonal guides targeting the same genes

As a metric for iMAP performance, we designed 2-3 guides per gene for four of the 70 RNA modification genes, totaling 10 guides for 4 genes (Figure S3A). These guides are predicted to produce null alleles, and the orthogonal guides indeed behaved similarly across all 46 tissues surveyed, albeit with a minor discrepancy: for the two guides targeting *Ythdf1* (g17-18), g18 was not altered in the germ line (T42-46), while g17 mildly (<1.5x) enriched (Figure S3A). Thus, among 460 data points (10 guides x 46 tissues), only 5 (1 guide x 5 germ cell types) are inconsistent among the orthogonal guides, comprising only ∼1% of total data points. Of note, for 2 other genes (*Fto* and *Gon7*), two guides are designed for each, but the guides are predicted to produce distinct effects, which was indeed fulfilled (Figure S3B-C).

### Seeming inconsistencies with the literature

The iMAP data agree well with the literature, but with three exceptions. In each case, guide abundance was altered in iMAP mice despite little defects in the KO mice, but each discrepancy can be reconciled by considering cell competition unique to the mosaic iMAP mice.

First, g42-*Elp1* was depleted in the epithelium of large intestine whereas conditional *Elp3* KO in the intestinal epithelium fails to produce overt phenotypes (*40*). Perhaps the KO cells are intrinsically defective, but the defect can only be exposed in the mosaic iMAP mice where the KO cells tend to be surrounded by, and thus potentially in competition with, normal cells in the same niche. Indeed, due to cell competition, some mutant phenotypes can only be exposed in mosaic systems (*41*, *42*).

Second, whereas g15*-Trdmt1* was marginally enriched in the cerebellum, *Trdmt1* KO mice are rather healthy (MGI:1274787), suggesting that this expansion might occur only in the mosaic iMAP mice, where the *Trdmt1* gRNA might enable the cells to become a “super competitor” to outcompete the normal neighboring cells; such an advantage does not exist in KO mice where all the cells are equally competitive (*43*). Alternatively, the putative cerebellum expansion might indeed occur in KO mice, but is well tolerated. Of note, g62-*Trmu* was also enriched in the cerebellum, but its KO mice die between implantation and somite formation, making it impossible to infer its role in the brain (*44*). Importantly, the overall mild and severe phenotypes in *Trdmt1* and *Trmu* KO mice, respectively, align well with the severe depletion across diverse tissues of g62*-Trmu* but not g15*-Trdmt1* (Figure 3D). Furthermore, conditional *Trmu* KO in the liver results in spotty liver necrosis, consistent with g62-*Trmu* being depleted in the liver in iMAP mice (Figure 3D). Thus, the literature on *Trdmt1* and *Trmu* KO mice tends to support the iMAP data.

Finally, g27-*Qtrt1* was depleted in spermatogonia (Figure 4F), but a gene-trap cassette that eliminates a 48-aa region required for QTRT1 catalytic activity fails to compromise spermatogenesis, which again can be explained by invoking the mosaic nature of iMAP tissues, especially because the trapped allele still expresses the remaining 90% of QTRT1 (aa1–104 and 151-403) and presumably retains most of its non-enzymatic functions (*45*).

### iMAP apparently outperformed lentiviral CRISPR screens *in vivo*

Genome-wide lentiviral CRISPR screens in the brain (*16*) and liver (*11*) provide a rare opportunity to benchmark the classic platform against iMAP. iMAP-91 showed much higher recovery rate of essential genes than the viral screens (Figure 3D), with a similar trend in iMAP-100 (*18*)—the first evidence of iMAP’s superior performance. To investigate further, we compared ROC-AUC values in the liver (brain lentiviral data unavailable). Using non-essential genes as negatives, iMAP achieved an AUC of 0.80, surpassing the lentiviral screen at 1 guide/target (AUC 0.73) and matching it at 5 guides/target (AUC 0.78; Figure S6A, white bars). Strikingly, with non-targeting guides as negatives, iMAP substantially outperformed the lentiviral screen even at 5 guides/target (AUC 0.92 vs. 0.82), likely reflecting its tighter null distribution (Figure S6A, gray bars). Multiple guides conferred only marginal benefits for the lentiviral screen—particularly with non-targeting negatives (ΔAUC = +0.03 from 1 to 5 guides/target)—as anticipated given its high-quality library powered by Rule Set 2 (Figure S6A; compare 1 vs. 5 guides). To dissect the basis for the AUC differences, we examined log₂ fold changes for guides targeting shared early embryonic-essential genes (n=25 guides) and non-essential genes (n=23 guides), plus non-targeting guides (n=16 for iMAP; n=1,813 for lentiviral). Both platforms correctly ranked depletion by essentiality (essential > non-essential > non-targeting controls; Figure S6B), but iMAP showed ∼4-fold stronger essential-gene depletion (median |log₂FC| ≈ 1.37 vs. 0.32; Figure S6B, left panel). This indicates superior knockout efficiency in iMAP, likely because its gRNA expression is more robust and uniform than the classic screens where the viral vectors are randomly integrated and high-expressers not enriched. Regarding the non-targeting guides, although both platforms exhibited near-zero median log₂ fold changes (0.02), the adjacent values range—which captures the bulk of non-outlier data—was ∼3.7-fold narrower in iMAP (width 0.23; −0.19 to 0.04) than in the lentiviral screen (width 0.84; −0.43 to 0.41; Figure S6B, right panel), signifying lower guide-to-guide variability in iMAP. The above fold changes were averaged across biological replicates, precluding assessment of inter-replicate variability. To address this, we computed the maximum-to-minimum fold-change ratio for each non-targeting guide across replicate mice (ratio of 1 indicates identical values). Although medians were comparable (1.29 for iMAP vs. 1.40 for lentiviral), the adjacent values range was 2.5-fold narrower in iMAP (width 0.49; 1.13 to 1.62) than in the lentiviral screen (width 1.23; 1.03 to 2.26), revealing greater mouse-to-mouse variability in the latter (Figure S6C)—likely attributable to its per-mouse viral library delivery. Collectively, these findings indicate that iMAP generates stronger on-target signals (perhaps via robust gRNA expression) and lower noise (perhaps from avoiding viral delivery), underpinning its superior performance despite using one guide per target. Side-by-side comparisons with identical guides would be required to confirm these advantages.

### Complexity of RNA modification factors

Our study sheds light on context-dependent gene functions and also reveals their complexity. Two issues are particularly noteworthy, each concerning tRNA modifications, as detailed below.

The first issue involves tRNA wobble uridine (U34), whose modification (mcm^5^s^2^) is important for accurate translation of a subset of tRNAs(*30*). mcm^5^s^2^ comprises a methoxycarbonylmethyl group at C5 position (mcm^5^) introduced by ELP1-6 in conjunction with ALKBH8 and TRMT112, and a thiol group at C2 position (s^2^) subsequently introduced by URM1 in conjunction with CTU1/2, with mcm^5^ being a prerequisite for s^2^ (Figure 3F)(*46*). Of note, ELP1-6 forms the Elongator Complex, whose main function is U34 modification rather than RNA polymerase elongation (*47*). iMAP-91 targets six of the 11 factors in the modification pathway, namely *Elp1, Elp3, Ctu1*, *Urm1, Alkbh8* and *Trmt112*. *Elp1/3* KO would eliminate all four modifications (cm^5^, mcm^5^, mcm^5^s^2^, ncm^5^) present on 11 tRNAs, whereas *Urm1/Ctu1* KO only mcm^5^s^2^ present on three (Figure 3F). Thus, one might expect stronger phenotypes from *Elp1/3* KO. However, *Urm1/Ctu1* KO is each lethal (MGI:1915455, 2385277), just like *Elp1/3* KO (MGI: 1914544, 1921445). In agreement with this, g*Ctu1*/*Urm1* depletion was as severe as g*Elp1/3* across diverse tissues and even far more severe in spermatogonia, suggesting that *Ctu1*/*Urm1* have functions beyond mcm^5^s^2^ writing. It is also unusual that although *Alkbh8* is crucial for mcm^5^s^2^ just as *Urm1/Ctu1*, g16-*Alkbh8* was only weakly depleted, and *Alkbh8* KO mice are alive with at most weak defects despite the complete loss of mcm^5^s^2^ (*48*, *49*). This is perhaps because in *Alkbh8* KO mice, (n)cm5 becomes highly abundant, which might compensate for the loss of mcm^5^s^2^ (*49*). In apparent conflict with this speculation, g73-*Trmt112* was severely depleted contrary to g*Alkbh8*, even though *Trmt112* acts in concert with *Alkbh8* in writing mcm^5^. The data suggest that *Trmt112* has functions beyond mcm^5^ writing and predict that *Trmt112* KO, which has not been reported, would be lethal.

The second issue regards the KEOPS complex mediating the N6-threonylcarbamoyladenosine (t6A) modification at A37 of a subset of tRNAs(*35*). KEOPS consists of five subunits (LAGE3, TPRKB, OSGEP, TRP53RK and GON7) each necessary for efficient t6A modification, with mutations in human orthologs each lead to Galloway-Mowat syndrome (GAMOS) characterized by organ anomalies and early death (*50*). However, although *Tprkb* and *Osgep* KO lead to preweaning lethality reminiscent of GAMOS (MGI:1917036,1913496), *Lage3* KO mice are viable with only a mild defect (neutrophil number change) unrelated to GAMOS (MGI:6257568), dovetailing with g38-*Lage3* depletion being much less conspicuous than g70-*Tprkb*/g79-*Osgep* depletion. Thus, in contrast to man, murine *Lage3* might be dispensable for t6A, or t6A dispensable for survival, which underscores species-specificity of KEOPS function. Of note, our study also bears on *Trp53rk* and *Gon7*, whose KO have not been reported. Specifically, murine TRP53RK is encoded by two genes, *Trp53rka* and *Trp53rkb*. g47-*Trp53rkb* (but not g20-*Trp53rka*) clusters with the guides whose target gene KO is lethal, predicting that *Trp53rkb* (but not *Trp53rka*) KO would likely be lethal. For *Gon7*, two guides were designed: g85-*Gon7-1* inactivates *Gon7* (together with two other genes), whereas g29-*Gon7-2* deletes GON7 C-terminal tail (Figure S3C). Guide depletion was widespread for g85-*Gon7*-1 but restricted to spermatogonia for g*Gon7*-2. Thus, *Gon7*/related genes are essential for diverse tissues, but the GON7 C-terminus is largely dispensable except for spermatogonia, highlighting the unusual vulnerability of this cell type.

### Therapeutic targets

First, we demonstrated the potential of iMAP in identifying targets for improving human NK-based therapy. Although the actual candidates emerging from the current screen of 70 RNA modification factors are limited in both quantity (only one gene) and quality (moderate potentiation of NK-mediated killing), larger-scale screens might uncover better candidates, which in turn necessitates the improvement in iMAP throughput. Our preliminary data suggest that a 10x-increase in the throughput (from 100 to 1000 guides/mouse line) is feasible (not shown).

Second, our study, together with a previous report that global *Alkbh5* KO selectively impacts spermatogenesis (*36*), highlights *Qtrt1/2* and *Alkbh5* as appealing male contraceptive targets.

Finally, our study bears on targeted cancer therapy. Guides targeting mouse orthologs of human CEGs were in general severely and ubiquitously depleted as expected, but g34-*Trmt5* was a striking exception, being hardly affected in most tissues despite normal editing rate (∼70% indel) (Fig. 3D, 3G). This could reflect an evolutionary divergence in *Trmt5* function (i.e., essential for human but not mice), or more interestingly, a dichotomy between cancer and normal cells (i.e., cancer but not normal cells is addicted to *Trmt5*). In the latter scenario, *Trmt5* inhibitors could become cancer drugs that are broad-spectrum yet safe. Such drugs would be transformative; indeed, eight CEGs have been FDA-approved as cancer therapy targets, but all of them are indispensable for normal cells, and their inhibitors are all highly toxic and of limited applications (*51*).

### Limitations of the study

First, the false negative rates are acceptable but still much higher than false positive (1/10 vs 1/590 in iMAP-61), which can be minimized by guide pre-validation via high-throughput screening (*52*). Second, some tissues might be sensitive to Cas9-induced double-stranded breaks, leading to false positives. Guides targeting safe regions can be included to control for such genome toxicity in the future. Finally, the *91-guide* array was constructed years ago, covering all the mouse RNA modification factors known at the time, but the number has almost doubled since then, necessitating the creation of new iMAP lines to catch up with the rapid progress in the field.

## Acknowledgments

We thank Dr. Kristin A. Knouse for advice, Drs Yong Cang and Peixiang Ma/Guang Yang for reagents, the Molecular and Cell Biology Core Facility (MCBCF) and the Molecular Imaging Core Facility (MICF) at the School of Life Science and Technology, ShanghaiTech University for technical support. This study is supported by National Key R&D Program of China (#2021YFA1100800, to T.C.), Open Project of BGI-Shenzhen, Shenzhen 518000, China (#BGIRSZ20220001, to T.C.) and Shanghai Municipal Government Local University Development Grant (#23010503300 to T.C).

## Author contributions

T.C. conceived of and supervised the project. B.L. and Z.J. generated the *91-guide* line, Z.J. created the iMAP database, G.W., Z.J. and W. W. did the experiments and/or bioinformatics analysis with technical help from the remaining authors.

## Data and code availability

All original code has been deposited as an R package at GitHub and can be accessed using the following link https://github.com/Jing-Zhengyu/iMAP. Any additional information required to reanalyze the data reported in this paper is available from the lead contact upon request. The source data and the codes for their plotting, together with the plasmid maps, are available at https:// github.com/Jing-Zhengyu/Data-of-iMAP. The raw data for NGS is available at http://www.ncbi.nlm.nih.gov/bioproject/1328907, with the accession ID being PRJNA1328907.

## Methods and Protocols

### Mice

The *91-guide* array, the ensuing transgenic line and the iMAP-91 mice were generated essentially as described (*18*). Briefly, we designed candidate guides using CRISPick and selected those likely to induce out-of-frame indels using inDelphi. Guide sequences are listed in Table S1. The transgene was assembled using Golden-Gate cloning (the sequence available at https://github.com/Jing-Zhengyu/Data-of-iMAP) and randomly integrated into the genome via *piggyBac* transposition following microinjection of fertilized eggs. PCR primers for determining transgene integration sites, transgene integrity, U6 promoter methylation, genotypes and P0 guide expression are listed in Table S1. The founder line bearing the transgene inserted at Chr6:69964770(+) (coordinates based on Mouse mm39) was used for further studies. Animal experiments were approved by the Animal Ethics Committee at ShanghaiTech, and performed in accordance with the institutional guidelines.

### Perturbation Atlas and comparison with lentiviral screen in the liver

The atlas was constructed essentially as described (*18*). Briefly, postnatal day 10 (D10) iMAP pups (*91-guide; Cre; Cas9*) were given TAM at 0.2 g/kg daily for four days and sacrificed 3-4 months later. Diverse solid organs/tissues were collected and genomic DNA extracted before PCR-NGS profiling of P0 guide representation. Immune and germ cell subsets were isolated by electronic sorting, and genomic DNA equivalent to ∼30,000 cells were used for PCR-NGS; PCR primers are listed in Table S1. The reads were normalized, fold-change over control (*91-guide*; *Cre*) calculated and data visualized in heatmap using a home-made script package (https://github.com/Jing-Zhengyu/iMAP). Briefly, the reads counts of targeting guides in experimental and control mice are first normalized to the non-targeting guides in the respective groups before deriving the ratios (fold-changes) of the reads counts. To compare iMAP with the lentiviral screen in the liver reported by Keys and Knouse, we first performed blast search for∼6500 control gRNAs listed in Table S2 of the Keys and Knouse paper against the mouse genome, which led to the identification of 1813 “non-targeting guides” that do not match the mouse genome. We then obtained gRNA reads counts from Table S3 of that paper for these non-targeting guides and also for all the targeting guides, and calculated their fold-changes relative to the plasmid library using the same script as for iMAP. The data were presented in Figure S6.

### Analysis of gRNA efficiency in N2a cells

We co-transfected vectors expressing gRNA (150 ng) or Cas9 (350 ng) into the mouse neuroblastoma N2a cells cultured on 48-well plates. 72 h post-transfection, we extracted genomic DNA, amplified 500-900-bp regions encompassing the sites targeted by gRNAs using 50 ng genomic DNA as templates. The amplicons were then sequenced using the Sanger method and analyzed using ICE (https://www.synthego.com). PCR primers are listed in Table S1.

### Screens for potential therapeutic targets in tumor-infiltrating NK cells

MC38 mouse colon cancer cells, gift from Yong Cang (ShanghaiTech University), maintained in DMEM supplemented with 10% FBS and 1% penicillin-streptomycin, were detached using 0.25% trypsin, washed with PBS, resuspended at 1×10⁶ cells per 100 μl PBS and injected the 100ul subcutaneously into the dorsal flank of 5-8 weeks-old iMAP-91 mice. Tamoxifen was administered via oral gavage (0.6 g/kg) on the day of tumor inoculation. Tumor growth was monitored every 2 days using calipers, and volumes calculated as (L x W^2^)/2, where *L* and *W* represent the longest and perpendicular diameters, respectively. Tumors were excised when reaching 1–1.5 cm³, which took 18-25 days, minced and digested in RPMI-1640 containing 1 mg/ml collagenase IV (Sigma-Aldrich) and 0.1 mg/ml DNase I (Sigma-Aldrich) for 20–30 minutes at 37°C with gentle agitation. The digested samples were filtered through a 70-μm strainer and cells pelleted by centrifugation at 400 ×g for 5 minutes (4°C). To identify TNFa regulators, cells were first stained for CD45.2-BB700, CD3-PE-Cy7 and NK1.1-APC, fixed with BD Cytofix/Cytoperm™ for 20 minutes at 4°C, washed with BD Perm/Wash™ and stained with TNFa-PE. CD45.2⁺CD3⁻NK1.1⁺ NK cells were sorted into TNFa^high^ (top 30%) and TNFa^low/-^(bottom 30%) populations using a BD FACS Aria III and analyzed by PCR-NGS.

### Target validation in human NK cells

Human PBMCs (Rubei Biotechnology) were cultured in Corning ® Lymphocyte Serum-free Medium supplemented with 10% FBS, 1% penicillin-streptomycin and 200 U/ml recombinant IL-2 (Novoprotein). NK cells were expanded using an IL-21-based NK Expansion Kit (Zhongwin Biotech) with stimulation on days 0 and 7. NK cell purity (>90% CD3⁻CD56⁺) was confirmed by flow cytometry on day 10. K562 leukemia cells stably expressing firefly luciferase (gift from Guang Yang, ShanghaiTech) (*11*, *53*) were maintained in RPMI-1640 with 10% FBS and 1% penicillin-streptomycin under standard conditions. RNP was assembled by incubating 40 pm each of recombinant Cas9 (Sino Biological) and sgRNA or non-targeting control (Table S1) at 37°C for 15 min. RNP was mixed with 3×10⁶ NK cells resuspended in 20 μl P3 Primary Cell Solution (Lonza) before electroporation using a 4D-Nucleofector™ (program CM-137). 24 h post-transfection, successful editing was confirmed by PCR-Sanger sequencing, and NK (0.2×10⁶/well) were co-incubated with K562 (0.4×10⁶/well) cells for 4-6 h in 96-well plates in PBMC culture media. To assess K562 lysis, cells were washed with PBS, resuspended in media containing cell-permeable D-Luciferin potassium salt (150 μg/ml, Beyotime) and incubated at 37℃ for 5 min before measuring luminescence using an Infinite M200 Pro plate reader (Tecan) (*53*). The lysis rate was calculated as (1-X/N) %, where X is the luminescence intensity of K562 cells co-cultured with electroporated NK cells, and N is that of K562 cells never exposed to NK cells. CD107a, perforin and Granzyme B were detected by FACS, whereas TNFa secreted to the media by ELISA.

---

**Figure S1.**
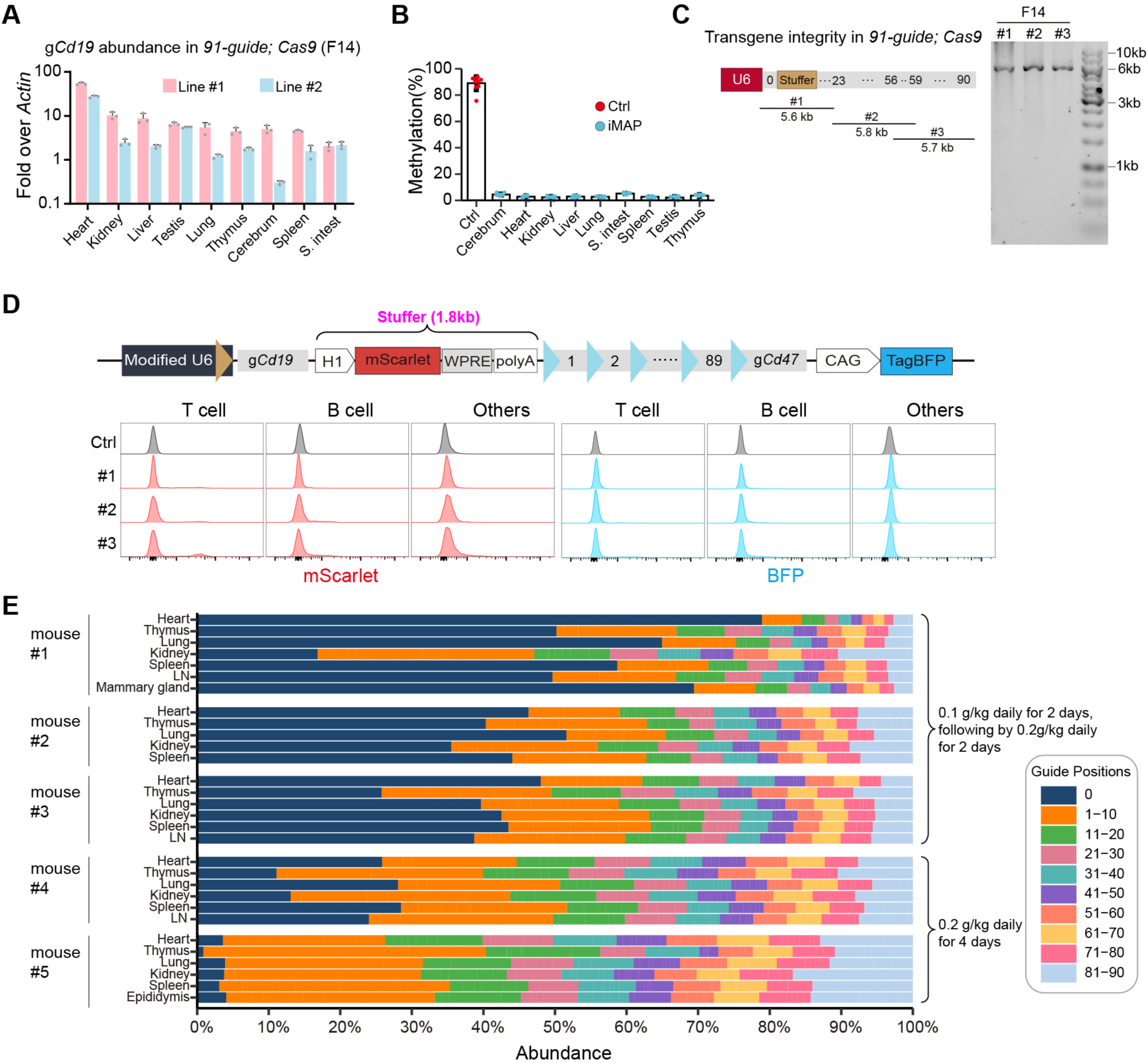
Basic features of the *91-guide* transgenic line, related to Figure 2 (A) qRT-PCR quantification of g*Cd19* expression in two transgenic lines. Line #1 was selected for further experiments. (B) Little U6 promoter CpG methylation at the U6 promoter. The *31-guide* transgene inserted into the H11 locus (red) serves as a positive control (*18*). (C) PCR analysis of transgene integrity. The three overlapping amplicons are diagrammed. The mice were *91-guide; Cas9* of the F14 generation. (D) FACS analysis of reporter protein expression in peripheral blood cells. mScarlet and BFP were expressed from the H1 promoter in the stuffer and the CAG promoter downstream of the *91-guide* transgene, respectively. The reporter genes were intended partly to facilitate the identification of the transgenic founder lines with transcriptionally active U6 promoter, as the transgene is randomly integrated via *piggyBac* transposition and thus subject to epigenetic silencing. Our previous study indicates that the CAG promoter activity cannot faithfully reflect the U6 promoter activity, suggesting the dichotomy in chromatin environment requirements for Pol II versus Pol III transcription (*18*). In this study, we added the H1 promoter as the second reporter, taking advantage of the fact that this promoter can direct both Pol II and Pol III transcription, and thus its Pol II promoter activity (manifested as mScarlet expression) might report the U6 promoter activity better than the CAG promoter. Unfortunately, this proved wrong: in the best founder line highly expressing gRNA as measured by RT-PCR (Figure S1A), neither mScarlet nor BFP expression was evident, reinforcing the notion about the dichotomy in chromatin environment requirements for Pol II versus Pol III promoters. (E) P0 guide composition of all the *91-guide; Cre* samples, a subset of them used in Figure 2A (bottom). Source data in Table S2.

**Figure S2.**
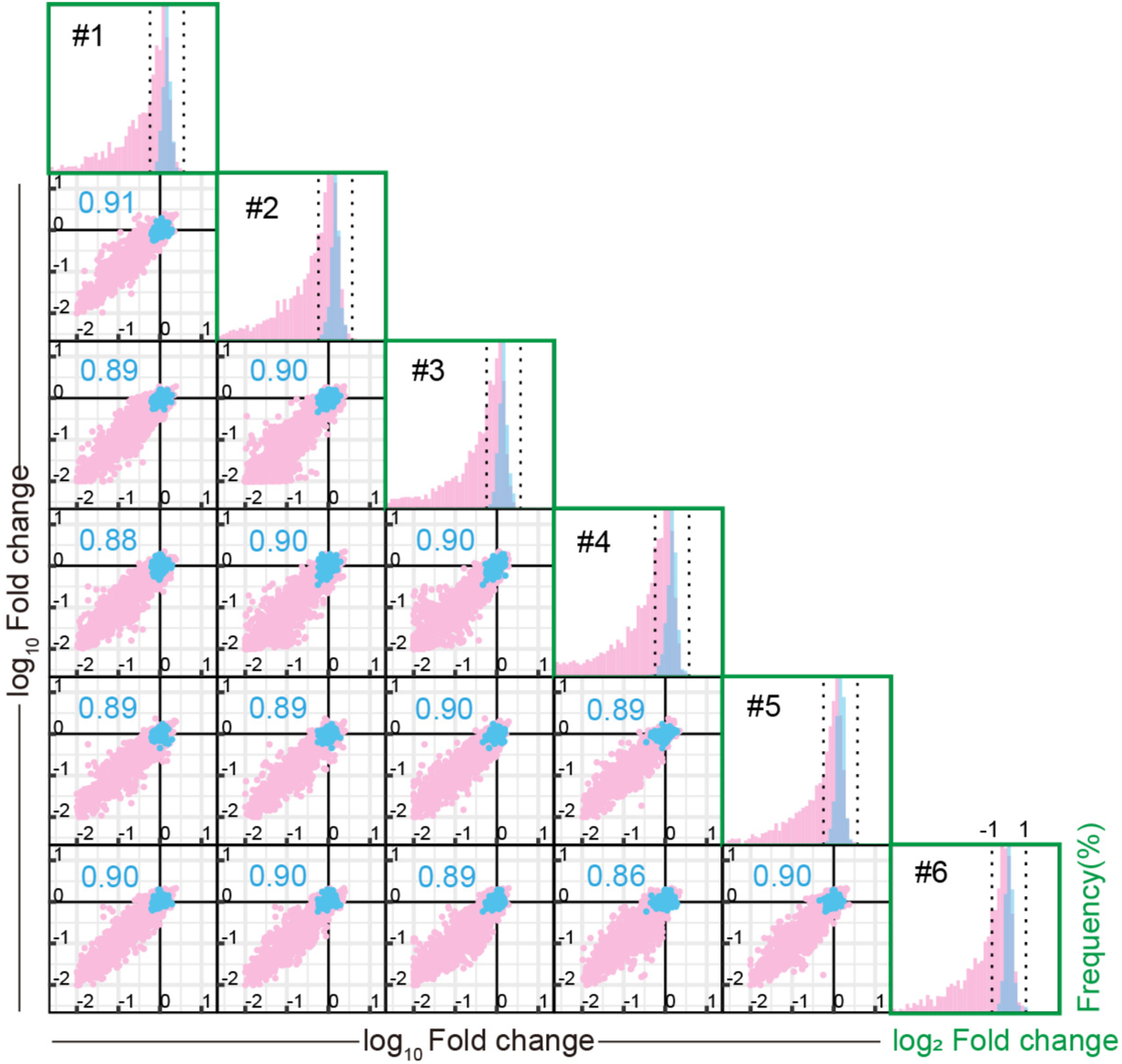
High replicate reproducibility of the P0 guide data in Figure 3B-D A total of 46 tissues, each with 2–6 (mostly 5-6) replicates, were sampled from six mice and fold-changes of each tissue replicate determined for each of the 87 guides listed in Figure 3D. The scatter plots display pairwise comparisons of the datapoints (tissue-guide combinations) among the six mice, together with the respective Spearman correlation coefficient. In the histograms, we binned the datapoints according to the fold-change magnitudes and then displayed the frequencies of each bin for each guide, with the highest frequency in each histogram set as 100%, as in Figure 3B. The fold-change magnitudes of the replicates and their frequency distribution patterns both prove highly reproducible.

**Figure S3.**
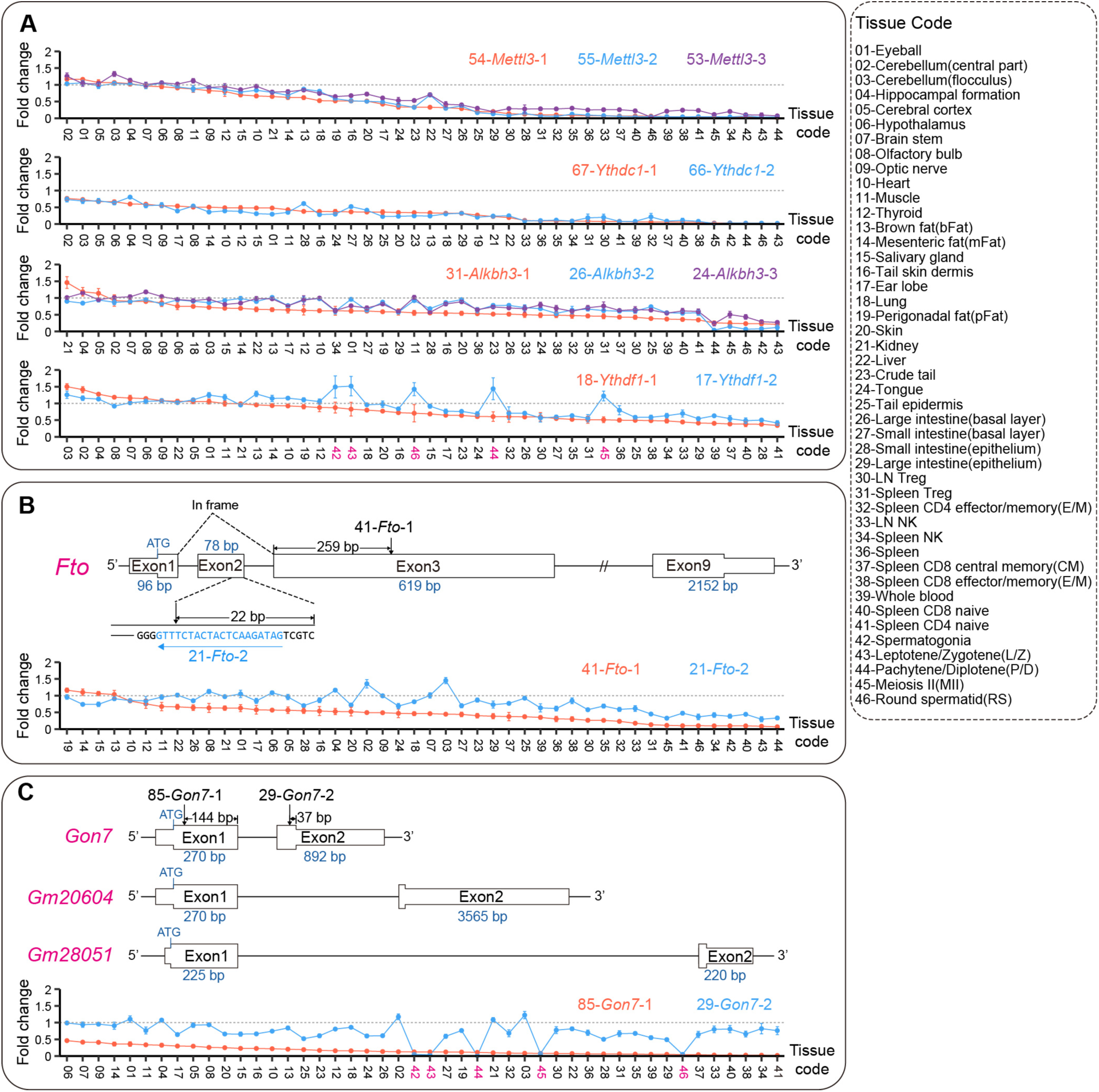
Analyzing different guides targeting the same genes, related to Figure 3D (A) Consistent depletion patterns of different guides designed to create null alleles of the same target genes. Of note, the two guides targeting *Ythdf1* (g17-18) closely co-segregated except that in the male germ cells (Tissue #42-46, pink), g17 (but not g18) was mildly enriched (<1.5x relative to NC). The discrepancy suggests off-target effects, reminiscent of the mild enrichment of g*Polr2a* in the testis in iMAP-61 line (*18*). (B) Distinct depletion patterns of guides creating hypomorphic vs null *Fto* allele. FTO is 502-aa protein encoded by 9 exons. g41-*Fto*-1 is predicted to truncate FTO at ∼aa128 to fully inactivate the protein, whereas g21-*Fto*-2 would likely cause Exon2-skipping by destroying its splicing donor, thus creating a partially active FTO mutant lacking the 26-aa fragment. (C) Distinct depletion patterns of guides creating hypomorphic vs null *Gon7* allele. GON7 is 98-aa protein encoded by two exons, the first shared with two related genes (*Gm20604*, *28051*). g85-*Gon7*-1 is predicted to fully inactivate *Gon7* together with the two related genes, whereas g29-*Gon7*-2 would likely create a partially active mutant lacking a∼13-aa tail. g29-*Gon7*-2 was selectively depleted in male germ cells (Tissue #42-46, pink).

**Figure S4.**
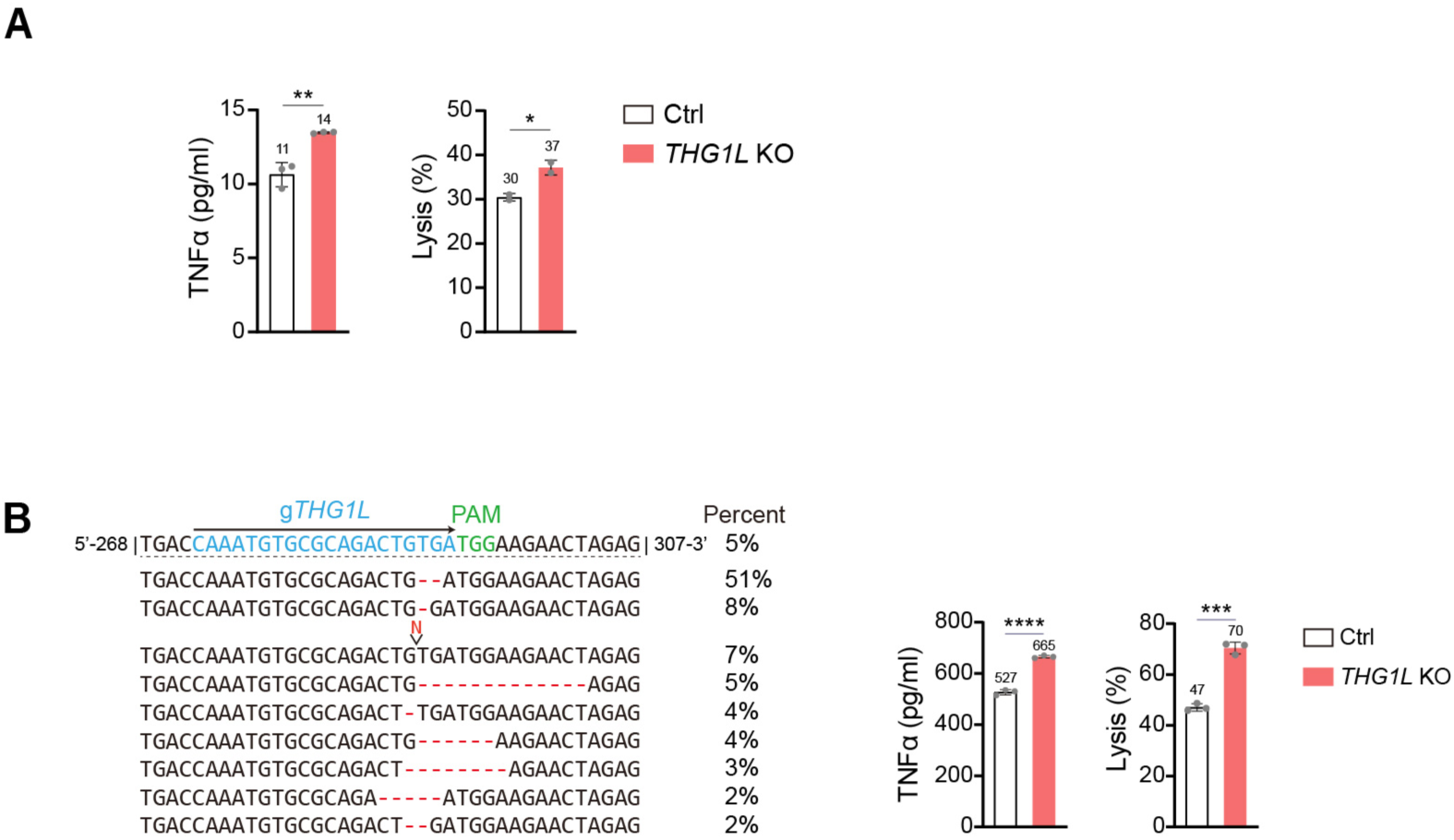
Target validation in human NK cells, related to Figure 5C (A) A biological replicate of the experiment in Figure 5C, confirming the effects of *THG1L* KO on human NK cells. NK cells were incubated with K562 tumor cells for 4-6 hr before analyses, as detailed in Figure 5B. The data again indicate that *THG1L* KO increased TNFα secretion concomitant with increased tumor lysis. Dots in the bar graphs represent technical repeats (n=2 to 3). *p < 0.05, **p < 0.01, Student’s t-test. (B) B is similar to Figure 5C, but orthogonal guides are used to target *THG1L*. Dots represent technical repeats (n=3). *p < 0.05, **p < 0.01, ***p< 0.001, ****p < 0.0001, Student’s t-test.

**Figure S5.**
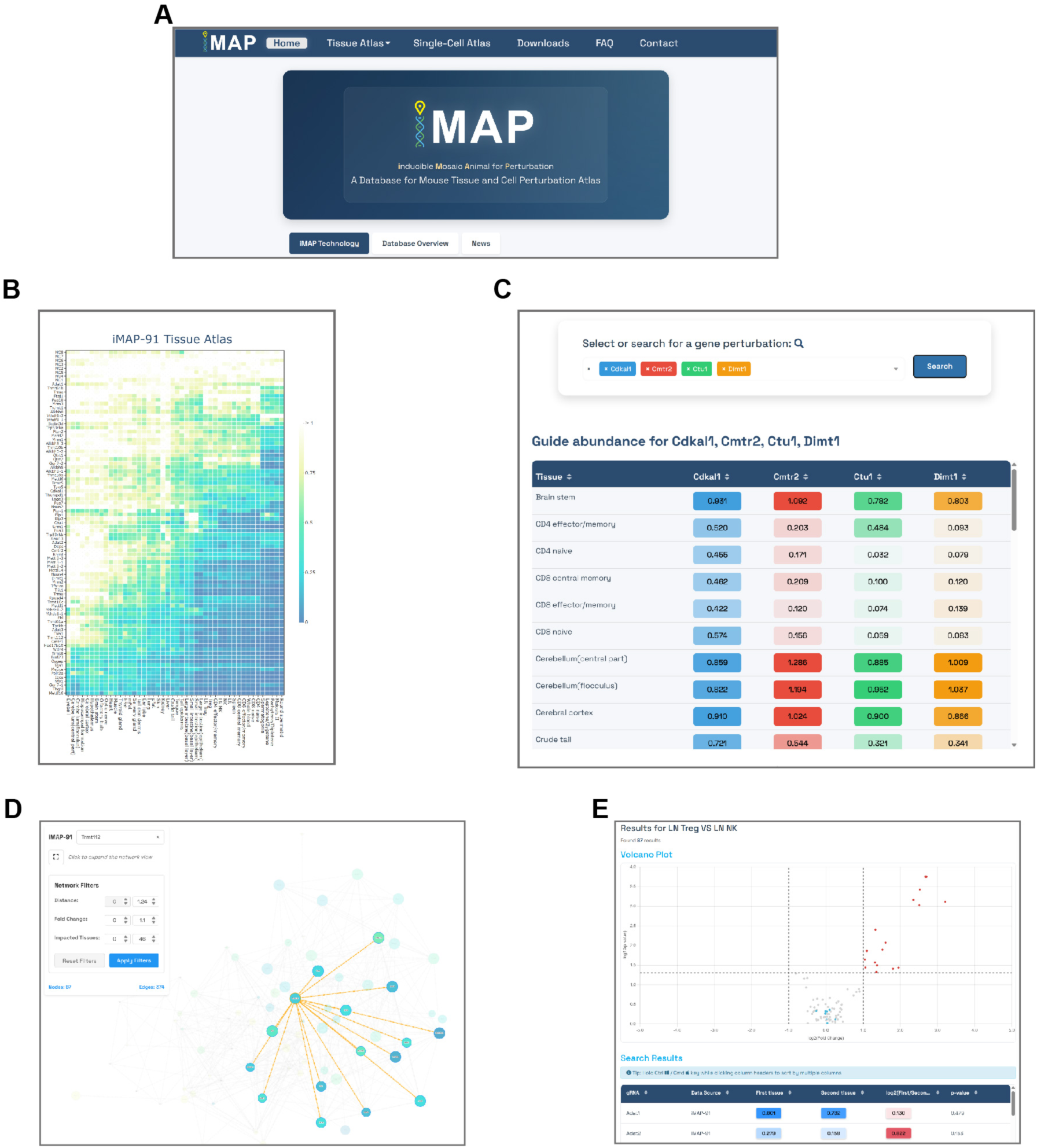
Snapshots of the iMAP database (A) Home page. (B) Heatmap view, displaying guide abundance across tissues. Actual fold-changes can be displayed by clicking on data points. One can also select a region of interest and zoom in. (C) Search result for the abundance of guides of interest across diverse tissues. (D) Search result for the network of guides surrounding a guide of interest. Each dot represents a guide. The color and size signify fold-change and the numbers of tissues affected, respectively. Additional information can be displayed by clicking on the guides and the lines connecting the guides. (E) Search result for differentially represented guides between a pair of tissues.

**Figure S6.**
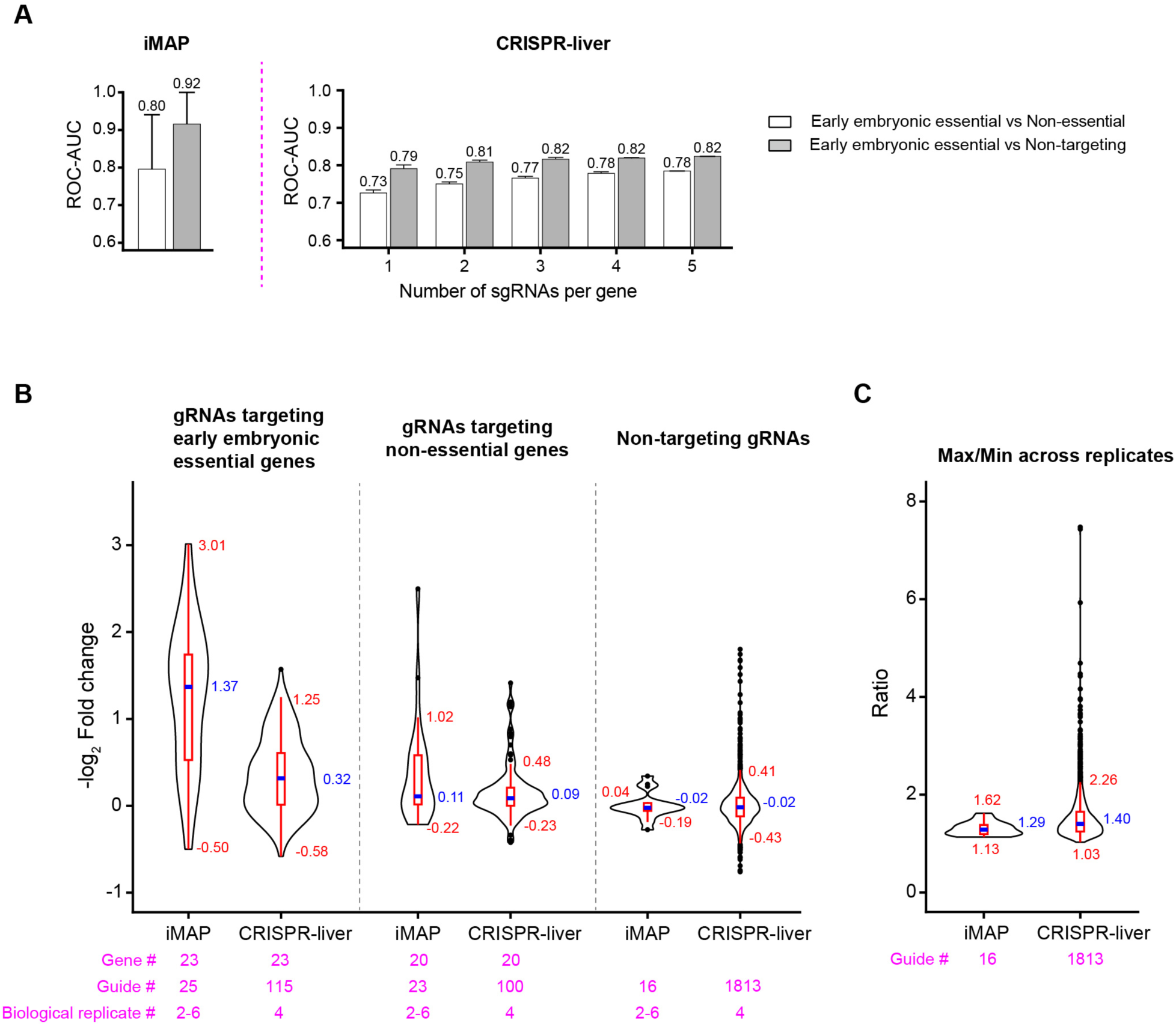
Comparing iMAP vs lentiviral screen in the liver (A) ROC-AUC analysis. (Left) iMAP. Data extracted from Figure 3H. Error bars represent 95% CI. (Right) CRISPR-liver screen. ROC-AUC is calculated based on subsampling of 1-5 sgRNAs per gene. Fold-changes of each guide are averaged from four biological replicates; error bars show s.d. across library subsampling iterations. The gold standard comprises early embryonic essential genes (n = 642) paired with either non-essential genes (n = 5006) or non-targeting guides (n = 1813). gRNAs were designed using Azimuth 2.0 (built on Rule Set 2), limiting the gains from additional gRNAs. Source data in Table S3. (B) Log₂ fold-changes for the targeting guides (left, middle) and non-targeting (right) guides. For genes with multiple guides, all the individual guides are shown. The iMAP data are pooled from iMAP-91 (6 replicates) and iMAP-100 (2 replicates) (*18*), while the lentiviral data (CRISPR-liver, 4 replicates) is from Keys and Knouse paper (*11*). Fold-changes of each guide are averaged from biological replicates as in A. The violin plots display mean fold-changes of each guide, with blue lines indicating the median, boxes representing the interquartile range (IQR, 25th to 75th percentiles), and whiskers defining the adjacent values extending from the quartiles to the most extreme data points lying within the Tukey inner fences (Q1 − 1.5 × IQR to Q3 + 1.5 × IQR). Points beyond these fences, if any, are shown as individual outliers. Source data in Table S3. (C) Mouse-to-mouse variability of non-targeting gRNAs. Fold-changes for non-targeting guides varied among biological replicates. To quantify this variability, the maximum abundance across the replicates is plotted relative to the minimum for each guide. The plot elements are as described in B. Source data in Table S

